# FLiTrak3D: Improved deep-learning-based 3D insect flight kinematics tracking using spatial and temporal encoding

**DOI:** 10.64898/2026.09.07.749842

**Authors:** Antoine Cribellier, Abel-John Buchner

## Abstract

Quantitative measurements of insect flight behaviour are essential for understanding the biomechanics, control, and ecology of flight, yet obtaining such measurements under free-flight conditions remains challenging. Small body size, rapid wing motion, visual symmetry, and frequent occlusions complicate three-dimensional pose estimation, often requiring restrictive experimental setups or substantial manual annotation.

We present *FLiTrak3D*, an open-source Python package for estimating insect flight kinematics from multi-view videography. It combines machine-learning-based markerless tracking with biomechanical modelling to reconstruct and parametrise insect body and wing motion. The workflow integrates image preprocessing, including dynamic image cropping and enhancement, two-dimensional bodypart localisation, three-dimensional reconstruction, and optimisation-based skeletal fitting.

A key innovation is the use of spatio-temporal encoding across synchronised camera views and adjacent frames to improve neural-network awareness of spatial and temporal context during bodypart localisation. Multi-view image stitching allows the network to use cross-view spatial relationships, reducing left–right bodypart misidentifications, while temporal encoding stacks consecutive greyscale frames into RGB images provide short-term motion information.

A species-specific skeleton is then fitted to reconstructed keypoints to enforce kinematic constraints, and estimate body and wing orientations. We demonstrate the approach using free-flying *Aedes aegypti* mosquitoes, achieving bodypart localisation errors close to human labelling, and realistic flight kinematics.

## 1 Introduction

Tracking the flight kinematics of insects is essential for understanding of their physiology and behaviour. Specifically, observations of insect flight kinematics can yield insight into the energetics of their flight [1, 2], their manoeuvring strategies [3], or their interactions, for example in swarms [4, 5, 6] or between prey and predator [7]. Linking kinematics with unsteady aerodynamic mechanisms helps illuminate evolutionary adaptation and ecological function[7], improve methods for the population control of problematic species such as disease vectors [8], and adapt insects’ locomotive strategies for bioinspired design [9]. For these reasons, insect flight has been a key subject of advances in animal pose estimation technologies. Yet, obtaining accurate 3D kinematics of flying insects remains challenging, and often requires substantial manual processing [10] and/or restrictive experimental setups [11].

Typically, pose estimation for animals, including insects, is performed via videography. Flying insects are particularly challenging to record because of their small size, fine-scale morphology, and rapid motion. One approach to simplifying the gathering of insect kinematics measurements involves physically tethering the insect to a fixed location. This circumvents these challenges, allowing detailed measurements within a restricted region, and enabling higher resolution and more controlled conditions. Consequently, tethering has been widely employed, especially in early studies (eg. [12, 13, 14]). However, kinematics measured from tethered insects risk being unrepresentative of their natural behaviour, due to the intrusive nature of tethering. Such data should thus be treated with caution [15, 16, 17], motivating the need for free-flight measurements.

Tracking insects’ kinematics in free flight is challenging, as it often requires analysis of sub-optimal image data; flying insects are often observed at low resolution because of the need to record over a large spatial volume relative to insect size, and images are sometimes blurry due to practical limitations on imaging depth-of-field. Various solutions have emerged to address this challenge, each with their own strengths and limitations. Attempts at automated kinematics extraction using conventional computer vision algorithms have frequently relied on the tracking of visible markers placed on the tracked animal (eg. [18, 19]). This can be impractical in insect flight experiments, due to small organism size, intrusiveness of the markers, or the procedure for applying said markers [11]. Automated kinematic extraction from videographic imagery without reliance on the application of markers has been achieved by, for example, Fry *et al.* [20, 21] and Liu and Sun [22], which manually fitted two-dimensional projected silhouettes to camera views. Such two-dimensional silhouettes have been widely used as a basis for three-dimensional hull reconstruction or shape carving methods, such as those developed by Ristroph *et al.* [23] and Walker *et al.* [24], the latter of which also allowed for the inclusion of wing twist, and was used in [25]. Such hull-carving algorithms are robust but suffer under conditions of (partial) optical occlusion [26] and require many cameras for an accurate shape reconstruction, while also struggling when significant morphological deformation is present [27]. Other algorithms have relied on highly specific experimental conditions [28] or been narrowly tailored to particular species’ anatomical structure or motion characteristics [29], limiting their applicability. Techniques for motion reconstruction from videographic imagery using traditional computer vision algorithms, as applied to insects and to animal locomotion more broadly, are reviewed by [30].

Where these methods require user input for each analysed frame, they are difficult to apply to the analysis of large datasets. This limitation has motivated the leveraging of powerful machine-learning algorithms, often based on deep neural networks (NN) [31], which have seen considerable development in recent years. Such machine-learning based approaches have demonstrated a scalability beyond that achieved by more traditional methods, due in part to a reduced requirement for manual human labour.

Two of the most popular 2D markerless pose estimators are DeepLabCut (DLC) [32] and LEAP/SLEAP [33, 34]. Both software packages exploit transfer learning to reduce labelling effort, and support multi-animal tracking [35, 36, 34]. Users of these packages can define animal bodyparts, label a limited number of images (*∼* 100 *−* 200 frames), and train a convolutional NN to automatically identify these bodyparts in similar images. Many related NN-based pose-estimation and tracking packages have since been developed, with varying levels of complexity, generality and support for single- or multi-animal tracking [37, 38, 39].

All these NN-based tracking packages fundamentally operate on two-dimensional data, requiring external processing tools for the reconstruction of the inherently three-dimensional motions of the observed bodyparts. Such a tool is 3DeepLabCut, which uses checkerboard-based calibration, provides multi-camera triangulation but does not impose articulated biomechanical constraints or estimate full kinematic parameters [40]. Similarly, the Python package AniPose [41], implements multi-camera triangulation with filtering and simple spatial constraints. Such triangulation relies primarily on geometric constraints and smoothing, without strong anatomical priors, so biologically implausible joint angles or temporal variation in anatomical dimensions may persist. Additionally, anatomical symmetry in visually similar bodyparts has the potential to cause ambiguities such as left–right bodypart swaps. In a similar vein, Günel *et al.* [42] developed DeepFly3D to synthesise multiple camera views of a tethered, walking, Drosophila into three-dimensional NN-based bodypart localisations. Uniquely, DeepFly3D mathematically relates the camera views via a self-calibration procedure based on the work of Chavdarova *et al.* [43], using the tracked insect as target. DeepFly3D further detects and removes erroneous bodypart localisations by checking cross-view consistency using the pictorial structure approach presented in [44], but this method did not eliminate the need for manual intervention.

A necessary step in understanding the behaviour of animals is the utilisation of tracked keypoints (bodyparts) for kinematic parameterisation. Current open-source 3D animal pose estimation tools primarily focus on the keypoint reconstruction aspect, with only limited support for downstream kinematic analysis. For example, AniPose [41] includes the possibility of computing joint angles from 3D keypoints, but more comprehensive estimation of kinematic parameters is not implemented in any of the surveyed packages. Additionally, none of the available toolkits provide functionality for fitting articulated 3D skeleton models to keypoint data, such as would allow for regularisation according to strong user-defined biomechanical constraints, and hence more accurate derivation of kinematic metrics [45].

An additional strategy for improving pose estimation accuracy is to incorporate spatial and temporal context directly into the neural-network inference stage. Standard markerless tracking frameworks typically analyse each camera view and frame independently, leaving the integration of information across cameras or timepoints to later processing steps such as triangulation or filtering. However, exploiting correlations across views or across adjacent frames during inference can help resolve ambiguities arising from occlusion, motion blur, or morphological symmetry. Several approaches have explored such ideas. For example, DANNCE performs multi-camera pose estimation by projecting synchronised camera views into a shared three-dimensional feature space and applying a 3D convolutional neural network to infer landmark locations directly from this combined representation. Temporal context has also been incorporated into pose estimation networks, for instance through architectures that process short frame sequences using convolutional or recurrent neural networks [46], or through extensions such as T-LEAP that explicitly integrate temporal information during inference [47]. While these approaches demonstrate the potential benefits of spatial or temporal context, they typically rely on specialised network architectures or training procedures. Methods that introduce such contextual information while remaining compatible with widely used two-dimensional pose-estimation frameworks would therefore provide a flexible and practical alternative.

We describe here an open-source Python package, *FLiTrak3D* [48], for estimating three-dimensional insect flight kinematics from videographic recordings from multiple simultaneous camera views. Our package constitutes a complete pipeline from raw videos to kinematic time series. It is designed to be adaptable to diverse insect geometries and explicitly addresses symmetry-induced ambiguities. The workflow consists of the following key processing steps: image preprocessing (enhancement, dynamic cropping, spatio-temporal encoding), machine-learning-based pose tracking in two dimensions via DeepLabCut [32], or other compatible 2D keypoint detectors, reconstruction of bodypart locations in three-dimensional space, and the fitting of a geometrically constrained insect skeleton model to estimate body and wing positions and orientations. A central contribution is our newly introduced image-level spatio-temporal encoding (multi-view spatial mosaics and frame-stack temporal encoding) that improves 2D keypoint localisation, while maintaining compatibility with existing 2D tracking packages. We release *FLiTrak3D* together with example datasets and documentation at https://git.wur.nl/cribe001/flitrak3d [48] to facilitate adoption and extension.

We have already applied our tracking method to studying the free flight kinematics of mosquitoes during active escape manoeuvres [49]. Here we provide a full description of our tracking method as implemented in *FLiTrak3D* and demonstrate the use of its unique capabilities in the accurate three-dimensional tracking and measurement of the flight kinematics of the yellow fever mosquito, *Aedes aegypti*, in free flight.

## 2 Tracking method

Estimating the 3D flight kinematics of an insect species requires high-speed recordings of flying individuals. For typical insect flight applications, FLiTrak3D is intended for recordings with approximately 20 images per wingbeat, using a minimum of 3 synchronised cameras placed at an angle of approximately 90 degrees from one other to maximize recorded 3D information (Fig. 1a). Using at least one top or bottom view simplifies tracking, as both insect wings will most often be visible in such a view. Back-lighting is often convenient to record well-lit images, but this can come at the cost of reduced ability to discern orientation of the flying insect from individual views. We will see later how stitching images from multiple viewpoints prior to analysis in DeepLabCut (or similar tracking software) can help resolve this problem (Section 3.4).

**Figure 1:**
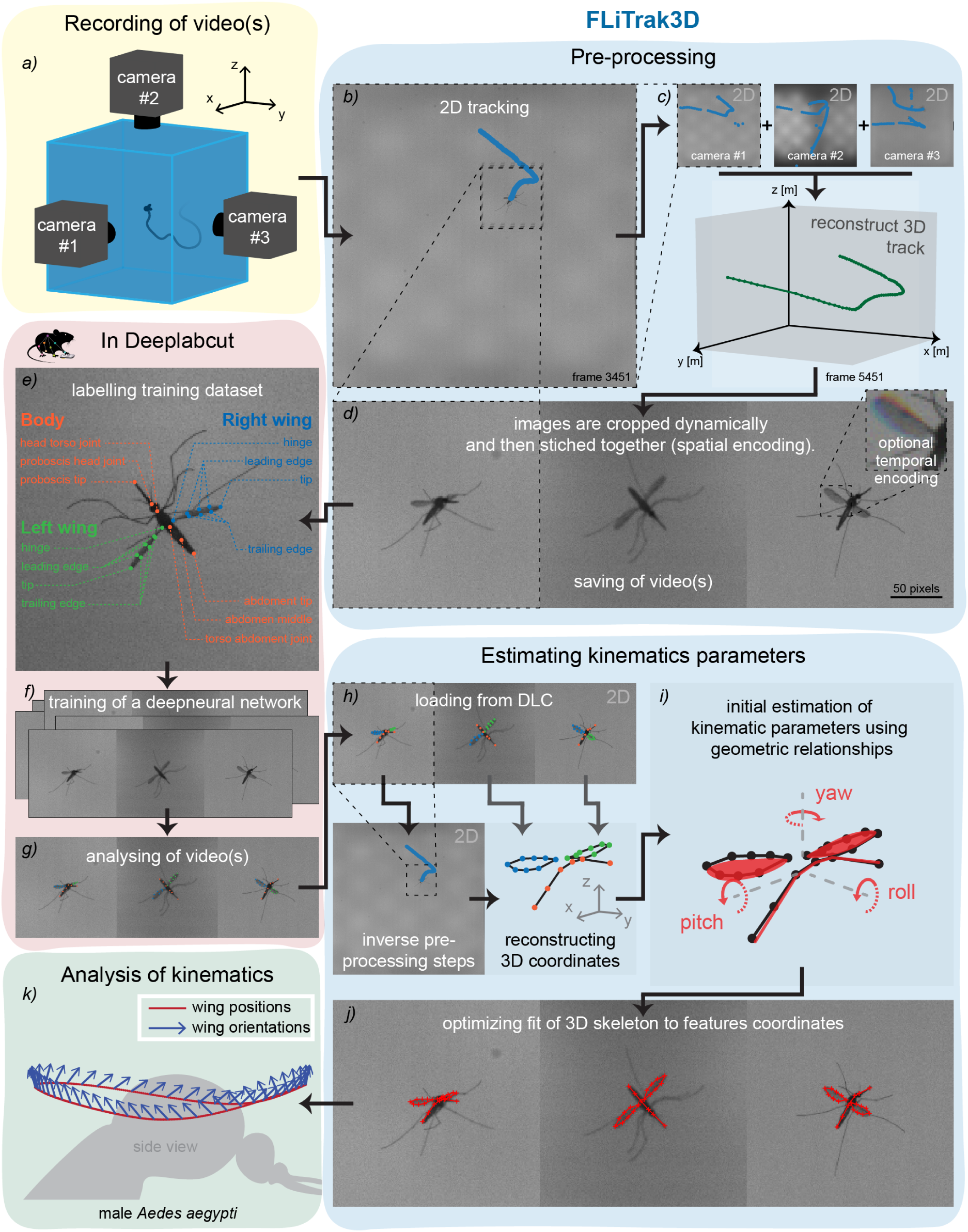
Flowchart of the tracking workflow. (a) Free-flying insects are recorded by cameras from multiple directions. (b,c) Two-dimensional detections are used to reconstruct three-dimensional trajectories. (d) Image regions surrounding the tracked insect are dynamically cropped and can be stitched into image mosaics (spatial encoding) and/or temporally encoded (see Fig. 6). (e,f) A subset of encoded images is manually labelled and used to train a DeepLabCut (or similar) neural network to identify user-defined bodyparts. (g,h) The trained network predicts bodypart locations in all frames, which are mapped back to the original camera views and reconstructed in three dimensions using the camera calibration. (i–k) A geometrically constrained skeleton model is fitted and optimised to reconstruct the insect pose and parametrise its kinematics. Data shown are from [49].

Camera recordings of a target insect’s motion can be automatically matched in *FLiTrak3D* across cameras by folder name or timestamp, while accounting for small timestamp mismatches across cameras. A calibration function relating each camera view to a global experimental frame of reference is defined. We used for this purpose a Direct Linear Transformation (DLT) model [50]. Pre-processing options include contrast/brightness/sharpness/colour enhancement, auto-contrast, histogram equalisation, view rotation, and optional resizing. Each applied action is reproducibly logged to a per-recording process history.yaml file.

To localise the animal before cropping, *FLiTrak3D* includes a simple background-subtraction and blob-detection-based 2D object tracking step (Fig. 1b), followed by a 3D reconstruction of these coarse tracks via the DLT calibration (Fig. 1c). This coarse 3D pass ensures the tracked object is consistent across views, and supplies centre locations in each frame for cropping. Frames are subsequently dynamically cropped to a region surrounding the moving insect to remove irrelevant pixels, reducing image size and improving computational efficiency..

The cropped views from all cameras are stitched together as mosaics to form single, combined frames, representing all simultaneous camera views (Fig. 1d), and saved to a new video file. The idea behind this stitching process is that it allows the NN, in subsequent training, to use simultaneous information from multiple camera views without requiring explicit knowledge of the multi-camera calibration. Because a single network processes the full stitched frame, corresponding bodyparts across views can provide contextual information that may help resolve visually ambiguous detections, particularly erroneous left–right bodypart swaps (see Section 3).

In addition to this spatial encoding (i.e. cross-view stitching), *FLiTrak3D* can perform temporal encoding by stacking the previous, current, and next greyscale frames into the RGB channels of a single image mosaic. Here we hypothesised that such temporal encoding supplies short-term motion context to the 2D keypoint detector, thus improving tracking accuracy without modifying the pose-estimation network architecture or requiring additional labels.

The next part of the workflow is performed within DeepLabCut (DLC) [32]. Note however, that the *FLiTrak3D* pre- and post-processing steps are tool-agnostic and can interface with other 2D keypoint detectors (e.g. LEAP/SLEAP [33, 34]) with minor adaptations. To allow subsequent kinematic extraction, the number and names of bodyparts labelled in DLC must be the same as defined in the selected skeleton in *FLiTrak3D* (e.g. see Fig. 1e for example labels for a two-winged insect). In DLC, bodyparts are defined by the user and located manually in a subset of frames, spanning the behaviours intended to be tracked and the diversity of the video recordings.

In support of achieving this goal, we recommend only labelling a few images per recording (i.e. *∼*3–5 frames if the insects do not change orientation significantly throughout the individual recording). These manually-located bodypart labels represent the ground truth location of each bodypart for use in training the NN (Fig. 1f). Typically for insect flight tracking applications we have found that a total of *∼*100–200 labelled images suffices. Still, we recommend only labelling *∼*50 images across *∼*20 recordings before training a first NN. This network should then be evaluated, to allow the user to assess the performance of the NN (e.g. via reprojection error), and to estimate where the NN still struggles to accurately track (e.g. for certain insect positions or orientations). Based on this assessment, new labelled images can be added to the training/testing dataset (for example per set of *∼*20 images) to iteratively improve the NN performance. As a final practical note, we found it crucial to regularly audit labels for errors such as left–right side swaps or misordered labels (e.g. “wing hinge” instead of “leading edge 1”). Any such mistakes remaining in the training/testing dataset can significantly impact inference performance of the NN.

The output of the neural network is a per-keypoint confidence map that defines a probabilistic estimate of the locations of bodyparts in each image. This is used to localise the bodyparts in each two-dimensional stitched image frame. Once each user-defined bodypart is tracked in the two-dimensional stitched camera planes using DLC (Fig. 1g), *FLiTrak3D* loads the NN-generated 2D bodypart detections along with the per-recording pre-processing log, process history.yaml. This allows *FLiTrak3D* to unscramble keypoints to the correct camera views, reverse any pre-processing (un-stitch, de-rotate, un-crop), filter by keypoint likelihood (default 0.7), and reconstruct 3D keypoint locations using the user-defined calibration (Fig. 1h). Both 2D and 3D bodypart locations are then written to CSV files.

A key aspect of *FLiTrak3D* is the regularisation of detected keypoints by fitting of a user-defined modular articulated skeleton model that reflects the animal’s morphology (see Fig. 2). The skeleton specification comprises (i) the set of bodyparts to be tracked, (ii) their connectivity (segments/joints), and (iii) optional biomechanical constraints. Segment lengths are treated as constant over time, and hard bounds can be imposed on joint angles and other kinematic parameters that must be respected during optimisation. These constraints suppress unphysical solutions arising from spurious 2D detections or transient 3D reconstruction errors.

**Figure 2:**
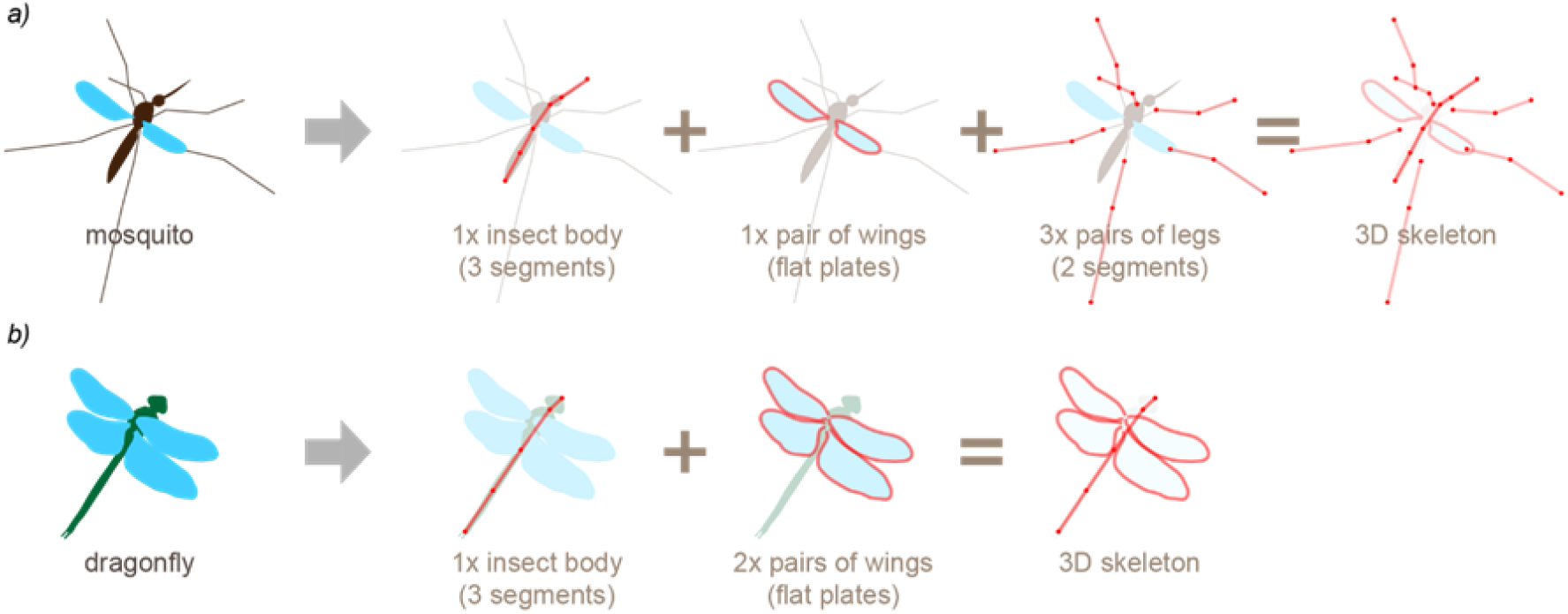
Example definitions of a modular insect skeleton model. (a) A two-winged insect (mosquito), including body, wings, and legs modules. (b) A four-winged insect (dragonfly) including only body and wings modules.

Fitting the skeleton yields a compact, interpretable parametrisation of the insect’s motion. The insect’s location in space is defined by the location of the thorax-abdomen joint in Cartesian coordinates (*x, y, z*), while its default orientation is given via the angle, *α*, of the thorax midline above the horizontal plane. Body orientation is further parametrised by yaw, pitch, and roll as extrinsic rotations about the fixed laboratory axes in the ‘xyz’ order, where the yaw is a rotation about the global *z*–axis (heading/yaw), the pitch about the global *y*–axis (nose-up positive), and the roll about the global *x*–axis; signs follow the right-hand rule. This ‘xyz’ extrinsic convention is equivalent to an intrinsic ‘ZYX’ Euler sequence (rotations about the body-fixed axes in the order *x*–*y*–*z*). Note that the pitch angle here is additive to the default thorax pitch orientation, *α*.

The position of each tracked wing is defined by three Euler angles: the stroke angle, *ϕ*, the deviation angle, *θ*, and the rotation angle, *ψ*, (Fig. 4d) as extrinsic rotations in the ‘yxz’ order from the yaw, pitch, roll rotated body coordinate system. The stroke angle, *ϕ*, is defined in the principal flapping axis, with its positive sense towards the insect’s anterior end. The deviation angle, *θ*, is defined relative to the stroke plane, with the positive in the upward direction for a mosquito in typical hovering orientation. A zero value of the rotation angle, *ψ*, corresponds to a wing leading edge oriented towards the insect’s anterior, with *ψ* = 90*^○^* indicating a vertical wing orientation with leading edge up, and *ψ* = 180*^○^* a wing leading edge oriented towards the insect’s posterior end.

An initial estimate of these parameters is made using simple geometric relationships (e.g. estimating stroke angle using wing hinge-to-wing tip orientation) (Fig. 1i). From these initial values, the articulated 3D skeleton model is optimised against the tracked keypoints (Fig. 1j) using user-selectable solvers (nelder-mead, powell, least squares, or leastsq from scipy.optimize), and loss is defined as the square root sum of squares of position errors. This can be done either directly in 3D (faster), where positions of the skeleton 3D keypoints are compared directly to those of the 3D keypoints reconstructed from DLC 2D estimations, or in 2D, where the skeleton 3D keypoints are projected to each 2D view and compared to original DLC 2D keypoint locations. The final output is a set of biologically interpretable kinematic time series, including body position, body orientation, and wing orientation angles, together with intermediate 2D and 3D keypoint data for inspection and further analysis (Fig. 1k). All processing steps are stored in a structured directory and can be reproduced non-interactively from the YAML log protocol process history.yaml.

## 3 Performance Evaluation

### 3.1 Experiment

To demonstrate our tracking method and quantify its uncertainty and limitations, we performed a free flight tracking experiment using *Aedes aegypti* mosquitoes. The mosquitoes (Rockefeller strain) were reared at the Laboratory of Entomology (Wageningen University & Research, NL), in 0.3 *×* 0.3 *×* 0.3 m cages (Bugdorm, MegaView Science, TW). The conditions under which the mosquitoes were raised included a constant temperature of 27*^○^*C and relative humidity of 70%. Concurrently, they were subjected to a 12 : 12 hour light:dark illumination cycle. The mosquitoes had continuous access to 6% glucose sugar/water solution and received a (human) blood meal (Sanquin, Nijmegen, NL) daily via a membrane feeding system (Hemotek, Discovery Workshop, UK). In these cages, female mosquitoes had access to wet filter papers upon which to lay their eggs. The eggs were collected and then dried for a period of 3 days before transfer to plastic larval trays filled with 27*^○^*C water containing a few drops of Liquifry No. 1 fish food (Interpet, UK). Emerging larvae were fed with TetraMin Baby (Tetra Ltd, UK). Pupae remained in their larvae trays covered with nylon netting material. Twice a week, emerged adults were vacuumed to new BugDorm cages. Males and females were kept together so they could mate.

Non-blood-fed adult mosquitoes of age 7.6 *±* 2.3 days (mean *±* standard deviation) after hatching were released into an octagonal, transparent acrylic flight arena of dimensions 50 *×* 50 *×* 48 cm (height *×* width *×* length) (described in detail in [49]). The arena was illuminated in the visible spectrum from above via an LED panel (OSLON SSL 80*^○^*, CS8PM1.PM, 20 *×* 48 cm), while 4 infrared LED panels (OSLON Black (850 nm) 150*^○^*, SFH 4716A) provided side illumination. The mosquitoes were recorded simultaneously from multiple directions using five Basler acA2040-90umNIR cameras fitted with Kowa LM12HC lenses (*f* = 12.5 mm, F1.4). Recording was performed at a framerate of 90 frames per second (fps), with camera sensors binned by a factor of three to 680 *×* 680 pixels. From these recordings, the mosquitoes’ three-dimensional point trajectories were tracked in real time using Flydra (v.0.20.30) [51]. A further three high-speed cameras’ (Photron SA-X2, 1024 *×* 1024 px) fields of view formed a volumetric sub-region of interest (ROI) of approximately 8 *×* 8 *×* 8 cm, close to the centre of the flight arena. A DLT calibration relates these three camera views to a shared global coordinate frame. The DLT coefficients were calculated from manually-located camera-plane locations of 26 spherical markers (diameter *d* = 3.5 mm) with known three-dimensional coordinates within the ROI. Upon detection (via Flydra-based tracking) of a mosquito entering the ROI, the three Photron cameras were triggered to each simultaneously record a series of 246 images at 12,500 fps. Recordings were taken in this manner over a period of several separate days.

### 3.2 Network definition and training procedure

#### 3.2.1 Experimental context and data

The dataset described in Section 3.1 presents a challenging tracking problem, although it is representative of other insect flight tracking settings. Bodyparts must be localised within greyscale, backlit recordings in which fine structural detail is largely suppressed: The silhouetted imaging conditions increase visual homogeneity across bodyparts. The small spatial scale of the tracked features, combined with a large recording volume and lighting constraints, results in partially focused, low-resolution images with relatively low signal-to-noise ratio. Accurate three-dimensional reconstruction further requires consistent tracking across multiple camera views, increasing sensitivity to optical occlusion and view-dependent ambiguities.

To reduce the size of the analysed data, images were dynamically cropped to 200 *×* 200 pixel regions centred on tracked reprojected 2D coordinates using simple 2D blob detection and 3D reconstruction (Fig. 1). Across 59 recordings, 295 sets of corresponding camera images were selected. In each camera view, 40 user-defined bodyparts representative of the anatomy of the mosquito (Fig. 1e) were manually labelled in DeepLabCut (DLC v2.3.0). Unless explicitly stated, DLC parameters were kept at default values in order to assess performance under typical usage conditions rather than pursuing dataset-specific hyperparameter optimisation.

#### 3.2.2 Data augmentation and training settings

Training data were augmented using the default imgaug augmenter [52]. Image scaling between 0.5*×* and 1.25*×* and rotations up to 25*^○^* were applied, but horizontal reflections and additional cropping were disabled to preserve geometric relationships between views, which are essential for cross-view learning. Pairwise prediction was enabled using a Huber loss [53] with weighting 0.1. Networks were trained on a single GPU for up to 100k iterations with batch size of 8.

A Part Affinity Field (PAF) model [54] was used to encode spatial relationships between bodyparts. Because full pairwise connectivity is infeasible for 40 bodyparts, a truncated PAF graph was defined manually to reflect anatomical connectivity. Within each view, the graph included body–leg connections and full connectivity among wing landmarks (80 edges per camera view). In stitched-view configurations, corresponding bodyparts across views were additionally linked (120 cross-view edges), resulting in 360 total connections (See supplementary Fig. S1).

#### 3.2.3 Train–test splits

To assess generalisation, two train–test regimes were defined:

- **Best-case (in-domain)** – For each of the 59 recordings, one randomly selected frame was withheld for testing (59 total test frames), while the remaining four frames per recording (236 frames total) were used for training. Test frames were unseen during optimisation, but originated from recordings represented in the training dataset. This ensures orientation and recording-specific visual features of the test dataset are well represented during training.
- **Challenge-case (out-of-domain)** – To assess generalisation to unseen recordings, eight entire videos (five frames each) were withheld for testing, and training was performed on the remaining labelled data. Test videos were selected to ensure approximately isotropic orientation coverage to avoid orientation bias. This split was repeated four times using different withheld recordings to assess repeatability. In this regime, the network cannot rely on recording-specific visual cues (e.g. minor lighting asymmetries or background artefacts) and must instead infer identity only based on learned spatial and cross-view relationships.

The selection of train and test data splits for the best case and challenge case are illustrated in Figure 3a and 3b, respectively.

**Figure 3:**
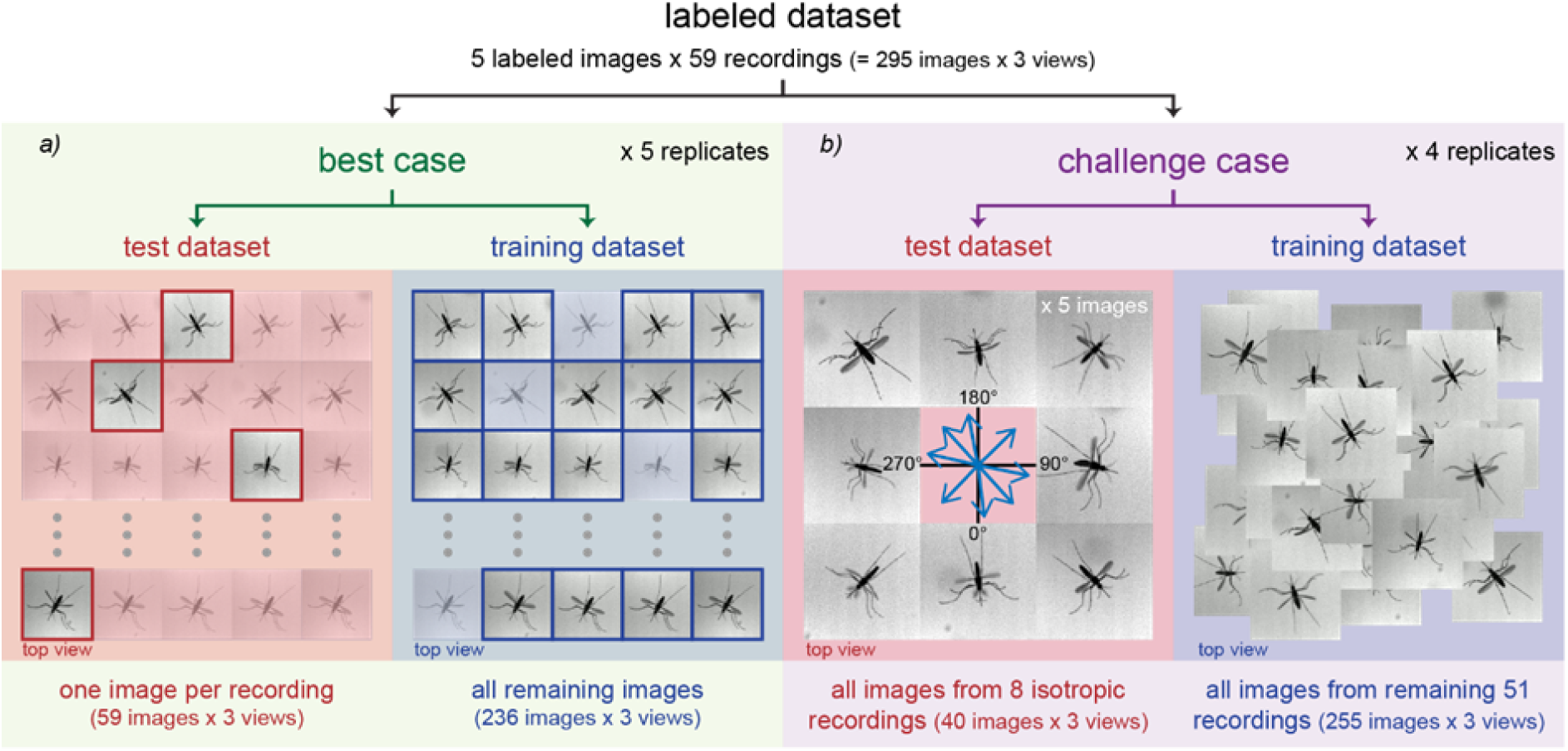
Definition of test cases for tracking performance assessment. (a) A best-case training draws a test dataset as a single labelled frame (unseen by network during training) from the same video recordings as the training dataset. Five independent replicates of this best-case training and network evaluation are performed. (b) A more challenging test case is defined by selecting eight from the 59 labelled video recordings as a test dataset, leaving the remaining 51 video recordings as the training dataset. The video recordings composing the test dataset are chosen such that the orientation of the flying insect is approximately isotropically distributed across the test dataset. Four independent replicates of this so-called “challenge-case” training and network evaluation are performed.

#### 3.2.4 Performance metrics

Prediction accuracy was quantified as the root mean square error *σ*, and median error *ɛ*, between predicted and manually-labelled positions. In each case, these metrics were averaged across multiple independently trained networks with differing train–test dataset splits. Because median error is less sensitive to body-part misidentifications, it is used as the primary accuracy metric, while keypoint misidentification rates are quantified separately.

Manual labelling uncertainty was estimated by re-labelling visible bodyparts in nine multi-view image mosaics (944 bodyparts per repetition), yielding *σ_m_* = 1.42 *±* 0.08 pixels (RMS *±* standard error of RMS) and *ɛ_m_* = 0.80 *±* 0.03 pixels (median *±* standard error of median). A bodypart was classified as correctly identified if the NN-predicted location lay within 3*σ_m_* of the manually-labelled location. An identification was considered incorrect if it failed this criterion and lay within this threshold of another manually-labelled bodypart location. A misassignment between bilaterally symmetric bodyparts was defined as a left–right swap (Figure 5d).

### 3.3 Best case performance

Under in-domain testing and using stitched image mosaics with temporal encoding, both RMS and median errors decreased rapidly over the first 10,000 training iterations (Fig. 4a-b). On the training dataset, RMS error fell below manual labelling variability at later iterations. On the test dataset, RMS error plateaued at approximately *σ ≈* 2 pixels (*σ* = 2.12 *±* 0.58 pixels at 100,000 iterations).

**Figure 4:**
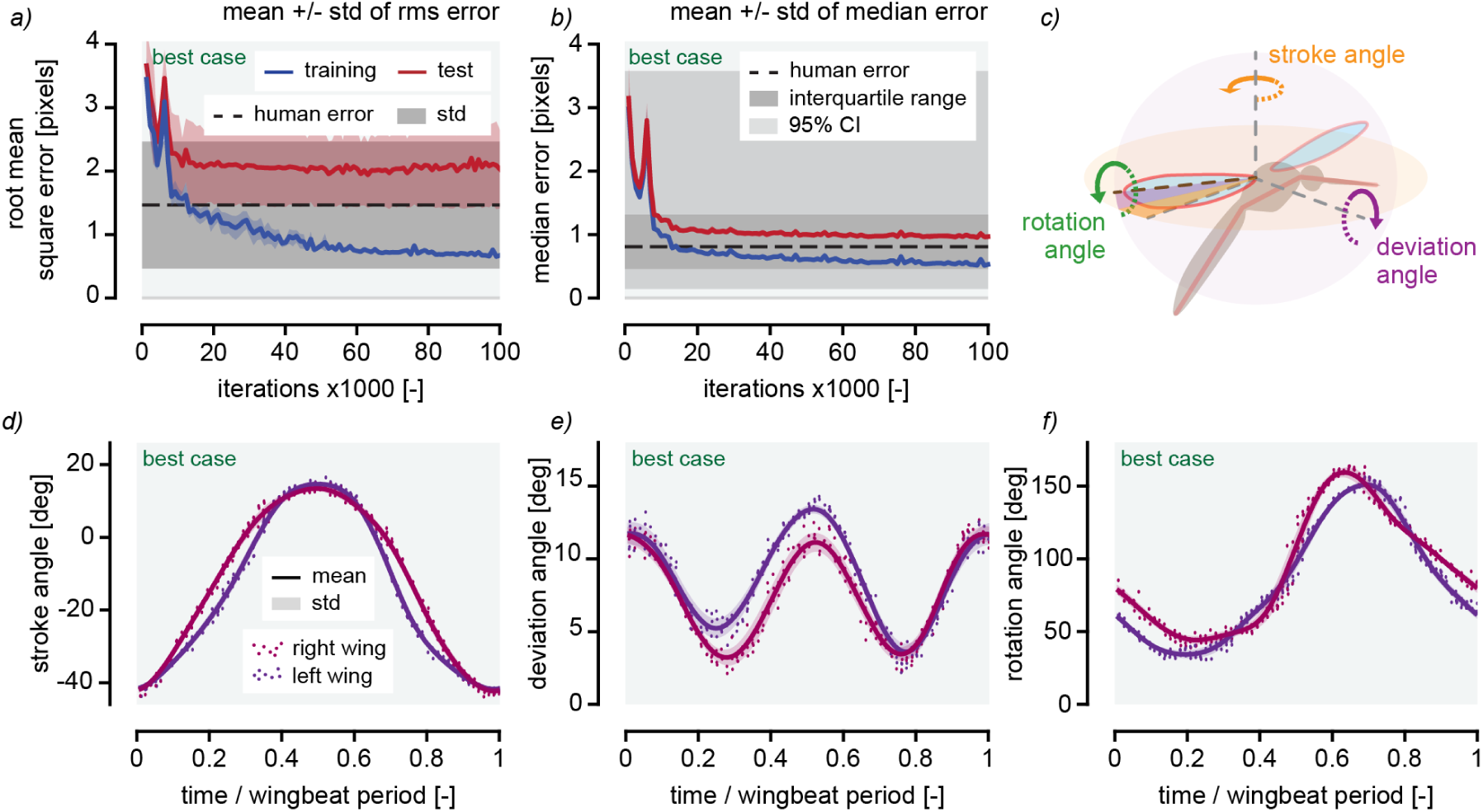
Tracking results for best case. (a) Root mean square error of bodypart localisation as a function of training iteration, for the best case network training. The blue curve represents evaluation using the training dataset, and the red curve represents evaluation using the test dataset. The black dashed line represents the root mean square human labelling error, as assessed via comparison of multiple independent re-labellings. Shaded areas represent the standard deviation across different bodyparts. (d) Median error of bodypart localisation as a function of training iteration, for the best case network training. Blue and red curves represent evaluation using the training and test datasets, respectively. The black dashed line represents the median human labelling error. Shaded areas represent percentile ranges across different bodyparts. (c) Orientation angles of a wing in space, defined as a stroke angle in the primary flapping plane, a deviation angle from that plane, and a rotation angle around the wing root–tip axis. (d-f) Wing orientation angles of a single automatically-tracked individual across a wingbeat period. Instantaneous angle identifications, tracked during 10 successive wingbeats, are given by blue and red markers, for left and right wings respectively. Solid curves represent the mean observed behaviour.

Median error similarly decreased rapidly and stabilised slightly above median manual labelling variability (*ɛ* = 0.97*±*0.05 pixels at 100,000 iterations). No increase in test error was observed at later iterations. The network correctly identified 86% of bodyparts with confidence ¿0.9, and left–right swaps were negligible (¡0.05%).

Three-dimensional reconstruction followed by skeleton fitting (section 2) produced wing Euler angle time series consistent with published mosquito flight kinematics (Fig. 4d-f) [25, 55]. In the example shown, an individual female beats her wings at a frequency of 528 Hz. Data tracked automatically from ten successive wingbeats in a single recording are plotted against their phase within a wingbeat period. A moving average (solid lines) indicates average behaviour of each wing over the included wingbeats. The wingbeat stroke amplitude in this example is approximately 55*^○^*, with a bias towards negative stroke angles (towards the mosquito posterior). The deviation angle varies at twice the fundamental wing beating frequency, between approximately 5-15*^○^* above the primary stroke plane which, in combination with the approximately-sinusoidal stroke angle variation, produces the figure-of-eight trajectory typical of mosquito wing-beating kinematics. The wing rotation angle ranges between 40-150*^○^*. The beating kinematics observed in this example differ slightly between left- and right-wings; small differences, which are consistently tracked by *FLiTrak3D* across the multiple wingbeats of the recording.

### 3.4 Challenge case performance

#### 3.4.1 The effect of cross-view stitching

In order to evaluate the effect of cross-view stitching (i.e. spatial encoding), three configurations were compared (Fig. 5):

- **Networks #1+**– Three independent single-view networks (one per camera) (Fig. 5a).
- **Network #2** – One single network trained on all single view images (Fig. 5b).
- **Network #3** – Stitched multi-view (spatial encoding) training with cross-view PAF connectivity (Fig. 5c, connectivity graph given in supplementary Fig. S1).

**Figure 5:**
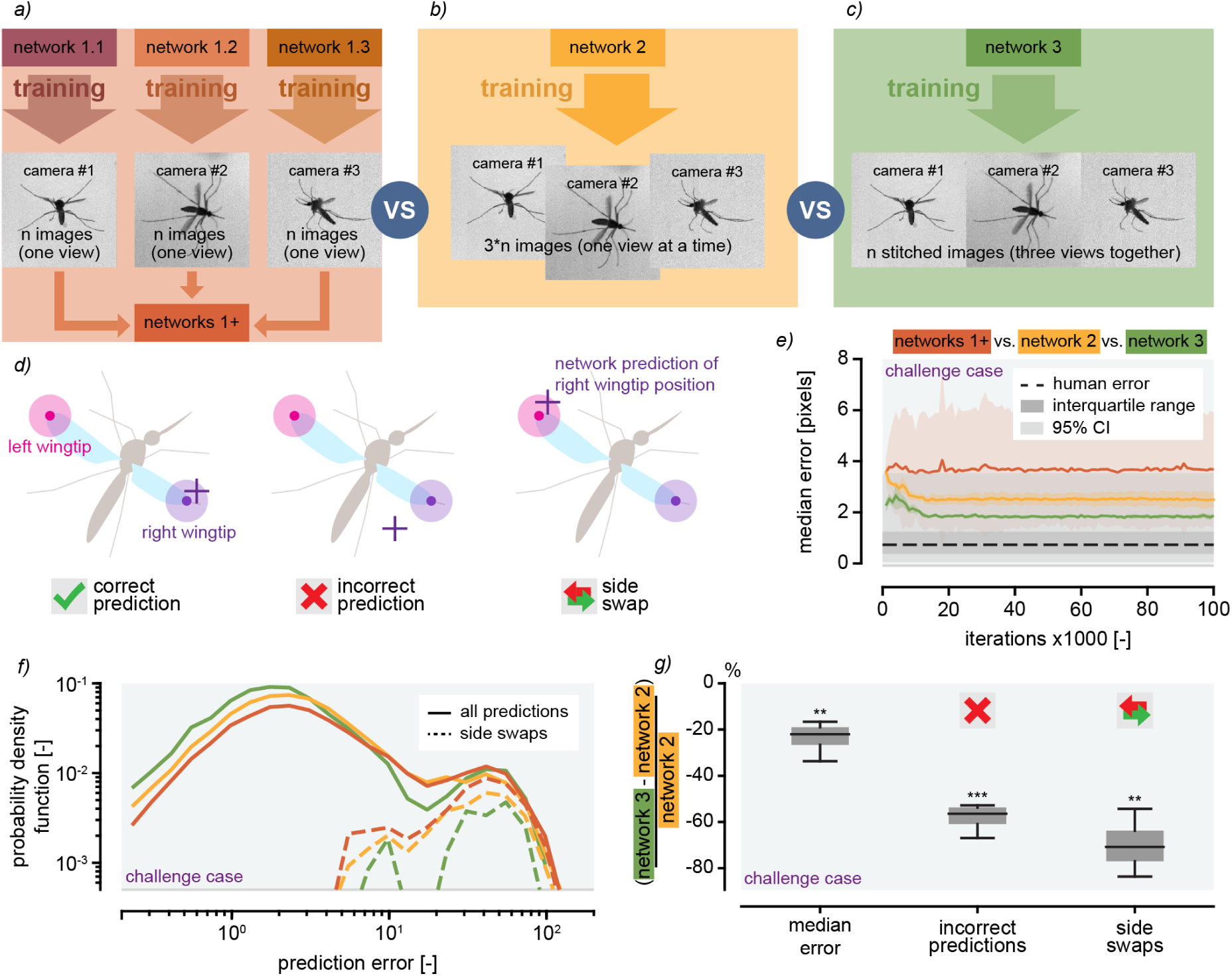
Setup and performance of the networks with the challenge case. Training data are organised in three different ways to test the effect of frame stitching on network training and performance. (a) Three separate networks are trained on unstitched camera views, each on views from a single camera (Networks #1.1, 1.2, 1.3), (b) network is trained on all labelled frames in all camera views, left unstitched (Network #2), and (c) camera views are stitched together, with bodyparts in each views defined separately and trained in a common network (Network #3). (d) Classification regime of network-derived bodypart localisations as correct, incorrect, or left–right swapped, using wingtip as an illustrative example. Light-shaded regions around bodypart locations represent three standard deviations of human labelling error. (e) Median error of bodypart localisations as a function of training iteration for all challenge case networks, at 100,000 training iterations. (f) Probability density function of prediction error for bodypart localisations for all challenge case networks. Prediction error distribution of localisations identified as side swaps are additionally plotted separately, under identical normalisation. (g) Comparison of median error, incorrect and side-swapped bodypart identifications by network #3 relative to network #2. Statistical significance of comparison with null hypothesis is shown using * (p¡0.05), ** (p¡0.01), and *** (p¡0.001).

Performance statistics from Networks #1+ are aggregated to enable direct comparison with Networks #2 and #3. Thus, each network of Networks #1+ are trained on n single images, while in Network #2, bodyparts from all views are pooled, effectively tripling the number of training images (3*n). In Network #3, each bodypart in each view is considered a separate entity for NN-training purposes, and therefore, the number of training examples per bodypart is the same as in Networks #1+, and one third as many as in Network #2.

All network configurations were trained to 100,000 iterations. Training and test images were not temporally encoded for this comparison. The median bodypart localisation error resulting from each network stabilised early, after approximately 10,000 iterations. Network #1+ performed, by this metric, the least well, with *ɛ* slightly less than 4 pixels (3.73*±*2.22, mean *±* standard deviation across replicates with differing train–test splits, Fig. 5e). The large uncertainty band reflects the variability in performance between networks #1.1, #1.2, and #1.3, which results from the differing viewing directions represented in each network’s training dataset. Inclusion of all camera views in the dataset upon which Network #2 is trained, improves performance such that *ɛ ≈* 2.5 pixels (2.53*±*0.36). Cross-view stitching in the construction of Network #3 further enhances accuracy to *ɛ ≈* 2 pixels (1.91*±*0.090). The error distributions (PDFs, Fig. 5f) produced by all networks are bimodal, with a primary peak near 1–2 pixels, and a secondary peak near 50 pixels; approximately the body length scale and thus likely composed of misidentifications. Error distributions of bodypart identifications classified as left–right swaps, according to the criteria illustrated in Figure 5d, reveal their predominance in this secondary peak and thus the mechanism by which many of these misidentifications occur. The error distribution of Networks #1+ exhibits the broadest primary peak and the highest rate of left–right swaps, while Network #2 shows reduced variance in the primary error peak, likely owing to its larger training dataset, but retains frequent left–right swapped misassignments. Finally, Network #3 yielded a narrower error distribution and substantially reduced left–right swaps. These performance improvements of Network #3 compared to Network #2 are quantified in Figure 5g. The accuracy, defined by median error, is improved by 23.6*±*7.45 % (one-sample t-test, p*<*0.01, N=4), but a larger effect is observed in reducing incorrect bodypart identifications by 58.2*±*6.41 % (p*<*0.001), of which side-swaps are reduced by 69.9*±*12.5 % (p*<*0.01). These improvements demonstrate the benefit of cross-view encoding for resolving bilateral ambiguity.

#### 3.4.2 Temporal encoding

We have demonstrated in the previous section that spatial encoding (i.e. cross-view learning) can have a beneficial effect in insect tracking, by reducing ambiguities in bodypart identification. Here we will explore if temporal encoding can similarly lead to an increase in tracking performance. To incorporate motion information, temporal context was encoded directly in the image representation. For a labelled greyscale frame at time *t_N_*, pixel intensities were stored in the green channel, while frames at times *t_N−_*_1_ and *t_N_*_+1_ were stored in the red and blue channels respectively (Fig. 6a). Such encoding implicitly provides directional motion cues (Fig. 6c) without altering network architecture or requiring further labelling.

**Figure 6:**
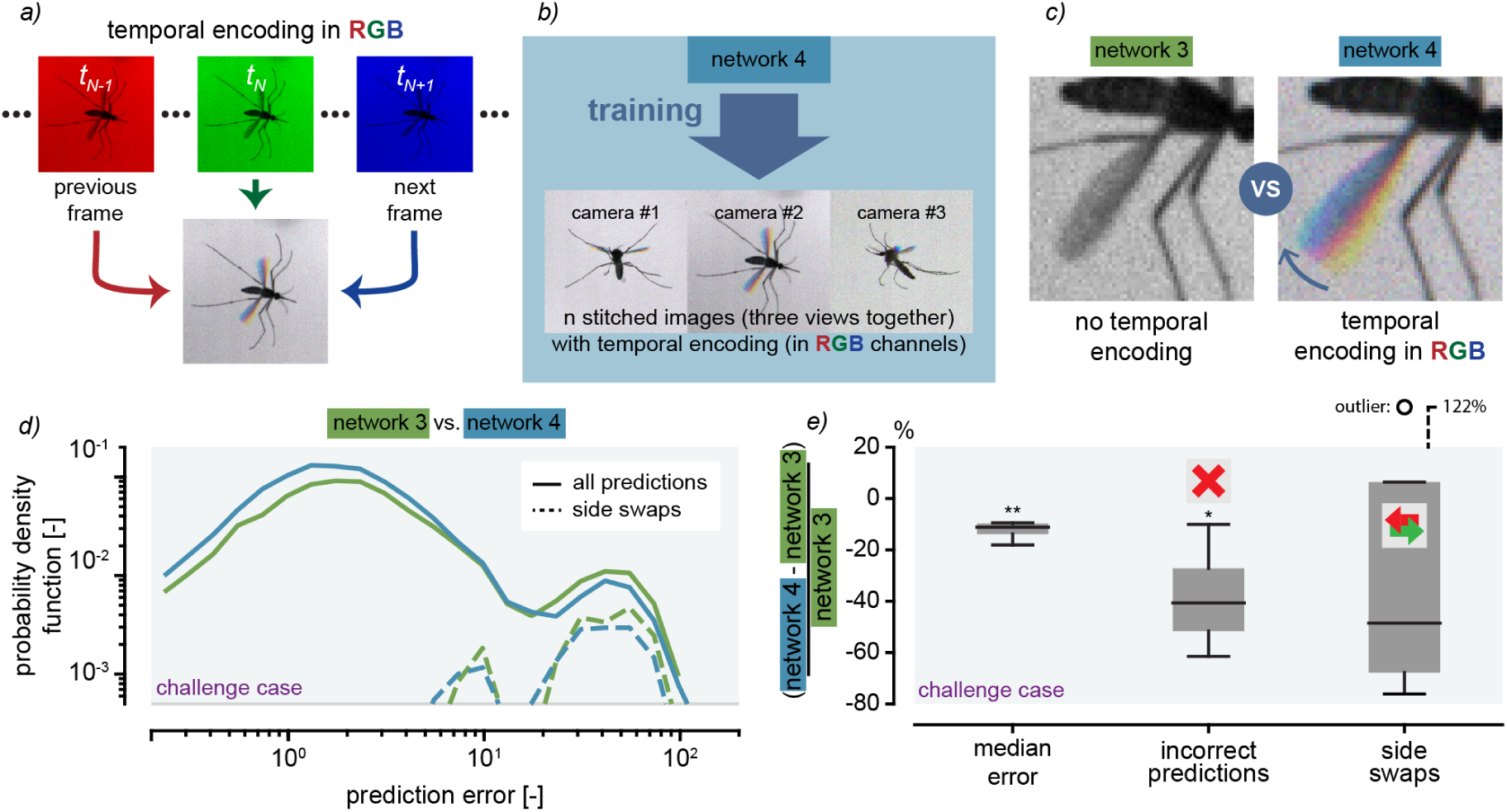
Setup and performance of the network using temporal encoding with the challenge case. (a) Encoding of temporal information is made using the black&white images at the previous, actual, and next frames to create a composite image with the pixel values of the three images in each of its RGB channels. (b) Like for the network #4, the RGB images (with temporal encoding) are stitched together, with bodyparts in each views defined separately and trained in a common network. (c) Comparison of images of a mosquito left wing with or without temporal encoding. (d) Probability density function of prediction error for bodypart localisations for the network with stitched images with (network #4) or without (network #3) temporal encoding on the challenge case, at 100,000 training iterations. Prediction error distribution of localisations identified as side swaps are additionally plotted separately, under identical normalisation. (e) Comparison of median error, incorrect and side-swapped bodypart identifications by network #4 relative to network #3. Statistical significance of comparison with null hypothesis is shown using * (p¡0.05), ** (p¡0.01), and *** (p¡0.001).

In this way, a spatially- and temporally-encoded network (Network #4) was trained with identical settings to Network #3 (Fig. 6b). The resulting error distribution remained bimodal, but the magnitude of the secondary peak was reduced (Fig. 6d). The primary peak increased in prominence and shifted toward lower error values. Median localisation error decreased across all test splits (mean reduction 12.5*±*3.97 %, one-sample t-test, p*<*0.01, N=4, Fig. 6e), and overall misidentifications were reduced by 38.3*±*22.1 % (p=0.0407). Left–right swap rates decreased on average, although the observed reduction (12.8*±*91.9 %, p=0.798) was not significant and one split showed an increase relative to Network #3. In that case, swap frequency remained below that observed in configurations without cross-view learning.

## 4 Discussion

Quantifying the three-dimensional flight kinematics of insects in free flight remains technically demanding. Small body size, rapid wing motion, optical occlusion, and limited visual detail, especially under backlit conditions, create conditions in which both classical computer-vision methods and standard two-dimensional pose-estimation pipelines face fundamental limitations [42]. Here we introduce *FLiTrak3D* [48] as a complete, open-source workflow that addresses these challenges from raw multi-camera recordings to biomechanically interpretable kinematic time series (Fig. 1). Beyond demonstrating tracking performance in *Aedes aegypti* mosquitoes, we show that integrating structured spatial and temporal data encoding into the pipeline substantially improves robustness in visually ambiguous conditions (Figs. 5 and 6).

Existing automatic approaches to three-dimensional insect flight reconstruction typically fall into two categories: Silhouette carving hull-based methods provide geometrically principled reconstructions but require many cameras and are sensitive to occlusion [26, 23, 27]. Neural-network–based pose estimators such as DeepLabCut and SLEAP offer scalability and reduced manual labour but operate fundamentally in two dimensions, requiring external triangulation and often lacking biomechanical regularisation [45]. Current 3D extensions primarily rely on geometric triangulation and smoothing, without explicit enforcement of realistically articulated anatomical structure. As a result, biologically implausible configurations or bodypart identity ambiguities, particularly left–right swaps, may persist.

*FLiTrak3D* addresses current methodological limitations across the full processing chain, from raw images to biomechanically meaningful kinematics (Fig. 1). Data preprocessing steps, which include dynamic cropping, centred on coarsely-tracked three-dimensional flight trajectories, image enhancement, and multi-camera calibration, are standardised and logged to ensure reproducibility across large and varied datasets. These preprocessing steps reduce the influence of background noise and decrease computational load, while preserving behavioural information. Spatial and temporal encoding can optionally be applied to these preprocessed images: stitching multiple camera views into combined mosaics allows a single network to learn cross-view correspondences directly from image data (Fig. 5), while leveraging the digital image colour channels for temporal encoding embeds short-term motion cues into the input representation (Fig. 6). Although cropping is recommended in most applications, all preprocessing and encoding steps remain modular and can be combined optimally on a case-by-case basis.

Preprocessed videos (e.g. .avi) can then be used directly with neural-network–based trackers such as DeepLabCut for 2D keypoint detection. Subsequently, *FLiTrak3D* loads tracked 2D image-plane points, automatically applies the inverse of the pre-processing transformation steps (e.g. un-cropping and un-stitching), reconstructs 3D keypoints in a calibration-defined global reference frame, and fits a user-defined articulated skeleton model with geometric and biomechanical constraints (Fig. 2). This final stage regularises noisy detections, enforces constant segment lengths and joint limits, and yields a compact parametrisation of body orientation and wing orientation angles suitable for aerodynamic and behavioural analysis (Fig. 4d-f).

We demonstrate the performance of *FLiTrak3D* on videographic data of free-flying *A. aegypti*, drawing training and test frames from the same recordings, and using both spatial and temporal encoding. In this best-case regime, the network achieved high accuracy (Fig. 4). Subtle recording-specific cues and a stable visual context likely reduce the effective complexity of the identity inference problem when testing on in-domain data, leading to near-human tracking performance (Fig. 4a-b).

In the challenge-case regime, where test frames originate from different recordings than the training frames, our performance analyses clarify the mechanisms by which spatial and temporal encoding improve tracking (Figs. 5 and 6). The principal failure mode of 2D keypoint detectors in this out-of-domain context is not small localisation noise but categorical misidentification of visually-ambiguous bodyparts, which is reflected in the bimodal error distributions we observe in the challenge case. The primary peak in the error distribution corresponds to near-human localisation precision, whereas the secondary peak reflects bodypart swaps at approximately body-length scale (Fig. 5f). Cross-view stitching (i.e. spatial encoding) substantially suppresses these large-magnitude identity errors, demonstrating that geometric redundancy across cameras resolves ambiguities introduced by bilateral symmetry and silhouette imaging (Fig. 5f).

Temporal encoding provides additional and complementary information (Fig. 6). By embedding consecutive frames into RGB colour channels, short-term motion direction becomes available to the network, allowing discrimination between symmetric structures such as leading and trailing wing edges. This exploits the fact of motion direction being a reliable distinguishing cue of bodypart morphology, due to physiological or behavioural constraints: for instance, the wing leading edge always precedes its trailing edge. Greater apparent motion in the image plane naturally enhances discrimination of motion direction, and thereby bodypart disambiguation. Temporal encoding consistently reduces median localisation error and incorrect predictions (Fig. 6d). Importantly, these gains are achieved without architectural modification, recurrent layers, or additional labelled data, and thus can be applied and tested on existing datasets, using existing NN-based tracking tools, with minimal effort.

Generalisation across recordings remains a critical issue for scalable behavioural analysis. In-domain validation can overestimate performance if networks exploit recording-specific cues. By evaluating out-of-domain train–test splits at the recording level, we show that spatial and temporal encoding enhances robustness to domain shift, an essential property for analysing large datasets in which labelling every recording is impractical. The ability of NN-based tracking tools to iteratively expand training datasets while monitoring reprojection errors, such as is facilitated by the *FLiTrak3D* workflow, further supports scalable deployment.

A further strength of the *FLiTrak3D* tracking pipeline lies in articulated skeleton fitting. Unlike pure triangulation approaches, optimisation of a modular morphological model suppresses unphysical solutions arising from transient 2D errors. The resulting parametrisation of body position and orientation, plus wingbeat stroke, deviation, and rotation angles, directly links tracking output to aerodynamic theory and behavioural interpretation. Thus, *FLiTrak3D* bridges the gap between pose estimation and kinematic analysis, in an open-source, user-friendly, comprehensive, and consistently documented analysis pipeline. Several limitations remain. Cross-view stitching presumes accurate temporal synchronisation and calibration, and errors at this stage may propagate downstream. Temporal encoding assumes locally smooth motion and may be insufficient for tracking during abrupt manoeuvres or extended occlusions, or in cases where biomechanical constraints on motion properties are relatively loose. Skeleton fitting requires species-specific model definition and careful parameter bounds, the choice of which necessitates prior anatomical and behavioural knowledge. Nevertheless, these constraints are transparent and user-controlled, and the modular design of *FLiTrak3D* facilitates adaptation to other taxa, including multi-winged insects for which the package already provides example modular skeletons (Fig. 2b).

In this work, we have presented *FLiTrak3D*, a package which provides an end-to-end, reproducible framework for extracting biologically meaningful 3D kinematics from multi-camera recordings of free-flying insects. Our results demonstrate that explicitly encoding spatial and temporal structure into the input representation, combined with articulated skeletal regularisation, substantially improves robustness in challenging visual regimes. More broadly, this work highlights the importance of representation and physical constraints in deep learning–based animal tracking, offering a scalable strategy for high-resolution behavioural quantification in complex biological systems.

### 4.0.1 Opening up

From a biological perspective, the ability to obtain robust, high-resolution three-dimensional kinematics in free flight expands the experimental space in which hypotheses can be tested quantitatively. In biomechanics and aerodynamics, it enables systematic testing of hypotheses regarding force production, flight energy demands, stability, and control across taxa and environmental conditions. In neuroethology, such data provides access how sensory cues are translated into flight manoeuvres during ecologically-relevant behaviours such as escape or host-seeking, where rapid manoeuvres are central. In behavioural ecology and collective behaviour, improved tracking of multiple individuals in shared volumes opens the possibility of resolving interaction rules in swarms or predator-prey encounters with greater precision. Beyond fundamental research, such measurements may inform vector-control strategies by linking flight behaviour to intervention outcomes, and support bioinspired robotics by providing empirically grounded templates for control in flapping-wing systems.

At a methodological level, our results highlight a complementary route for advancing machine-learning-based tracking: improving how information is structured rather than only increasing model complexity. By embedding spatial and temporal context directly into the input representation, and by enforcing biomechanical constraints during reconstruction, we show that robustness to visual ambiguity and domain shift can be improved without modifying network architectures or substantially increasing training data.

By manipulating the data representation at the input level to structurally include spatial and temporal context, and by enforcing biomechanical constraints during kinematic reconstruction, we demonstrate improved robustness to visual ambiguity and cross-recording domain shift. Crucially, these gains require neither modifications to network architecture nor substantial increases in training data volume, allowing application to existing datasets and workflows.

This challenges the prevailing emphasis on scale (larger datasets, deeper networks) as the primary driver of generalisation, and instead points toward hybrid strategies that integrate data-driven inference with domain-specific knowledge. Such an approach is likely to be broadly applicable in biological and biomedical settings where observations are noisy, incomplete, and governed by underlying physical constraints, suggesting our work can contribute towards developing more general frameworks for extracting reliable quantitative information from complex living systems.

## Ethic

This work did not require ethical approval from a human subject or animal welfare committee.

## Data Access

Code, examples, and documentation for *FLiTrak3D*, are publicly available at the git repository https://git.wur.nl/cribe001/flitrak3d [48]. Further supplementary material is available online with this article.

## Authors contributions

A. Cribellier conceived the original study, developed the software and methodology, and acquired the experimental data. A-J. Buchner contributed to the study design and methodology, and did the data curation. He analysed the data, and generated comparison between conditions be as well as the statistical analysis. Both authors contributed to formal analysis, validation, and visualization. Both authors also drafted the original manuscript, reviewed it, approved the final version for publication, and agree to be accountable for all aspects of the work.

## GenAi useage

The authors used AI-based tools solely for editing and formatting of the manuscript. All scientific content, interpretations, and conclusions were developed by the authors, who carefully reviewed and verified the final text and take full responsibility for its accuracy and integrity.

## Acknowledgment

We thank Cees Voesenek for the significant support and advices for the development of *FliTrak3D* and related Python library *TrakCoreLib*. We thank Florian Muijres for the guidance and various discussions on insect flight tracking and aerodynamics. We thank Tessa Visser, Pieter Rouweler, Kimmy Reijngoudt, André Gidding, and Frans van Aggelen for rearing the mosquitoes in Wageningen.

