## Supplementary material for "FLiTrak3D: Improved deep-learning-based 3D insect flight kinematics tracking using spatial and temporal encoding": Suplmentary figures

Antoine Cribellier Abel-John Buchner

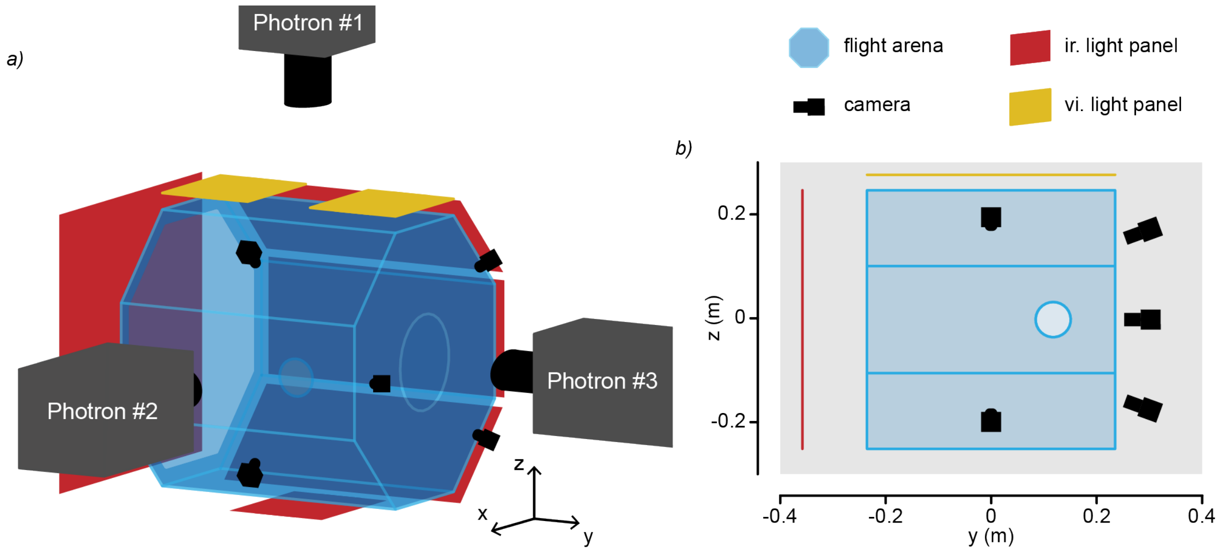


Figure S1: Experimental setup used to record mosquito flights. (a) three-dimensional and (b) two-dimensional views of the experimental setup showing the 5 small Basler cameras used to tracked mosquito in real time and that triggered the 3 large Photron cameras used to record high speed and high resolution videos of free flying *Aedes aegypti* mosquitoes.

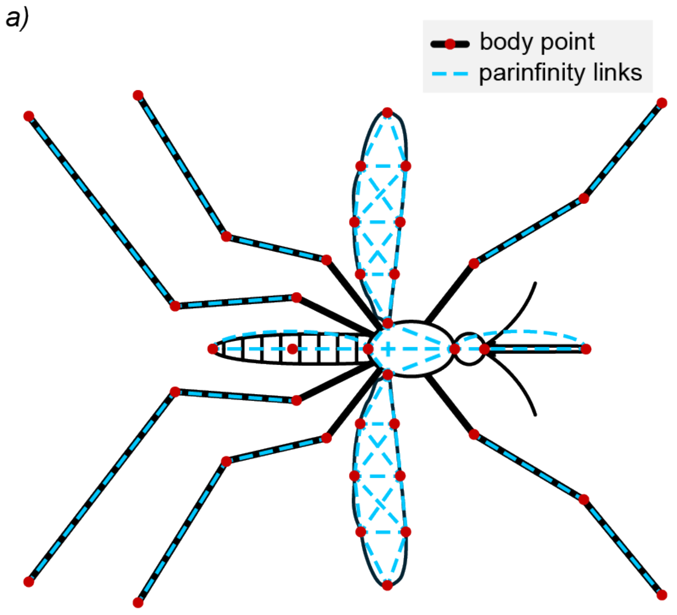


Figure S2: Part affinity field (PAF) graph connectivity within a single camera view. In Network #1, additional graph connections related each bodypart in each camera view with its counterpart in the complementary camera views.

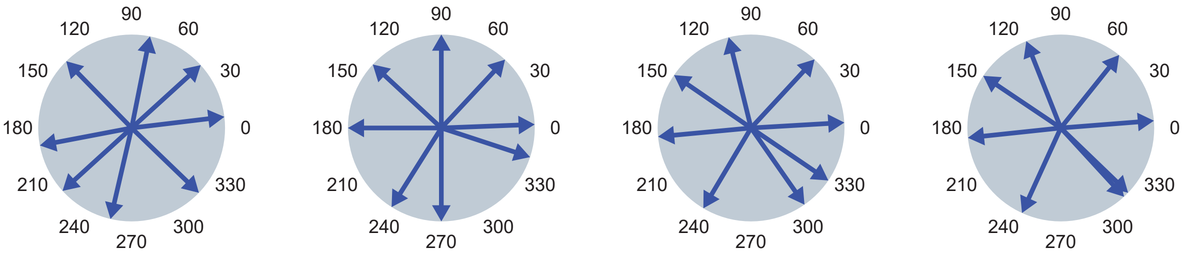


Test set #1 Test set #2 Test set #3 Test set #4

Figure S3: Mosquito orientation vectors of abdomen tip to proboscis tip. Orientations are in the horizontal plane, as viewed form the perspective of camera 2. Each set of 8 vectors relates to a single test–train split used in the challenge case evaluation. The combinations of orientations were chosen manually so that each sets encompass orientation in all directions

**Supplementary File S1:** **`config.yaml`.** DeepLabCut project configuration file used for training and evaluation of the spatially encoded multi-view mosquito tracking dataset. The file defines bodypart labels, skeleton connectivity, dataset settings, and training parameters.

**Supplementary File S2:** **`pose_cfg.yaml`.** DeepLabCut pose-estimation configuration file used for training the spatially encoded mosquito tracking network. The file specifies network architecture, augmentation settings, optimisation parameters, and the Part Affinity Field graph linking bodyparts within and across camera views.
